# Simultaneous Single-Cell Genome and Transcriptome Sequencing of Termite Hindgut Protists Reveals Metabolic and Evolutionary Traits of Their Endosymbionts

**DOI:** 10.1101/2020.12.11.422253

**Authors:** Michael E. Stephens, Jacquelynn Benjamino, Joerg Graf, Daniel J. Gage

## Abstract

Different protist species which colonize the hindguts of wood feeding *Reticulitermes* termites are associated with endosymbiotic bacteria belonging to the genus *Endomicrobium*. In this study, we focused on the endosymbionts of three protist species from *Reticulitermes flavipes*, which included *Pyrsonympha vertens*, *Trichonympha agilis*, and *Dinenympha* species II. Since these protist hosts represented members of difference taxa which colonize different niches within the hindguts of their termite hosts, we investigated if these differences translated to differential gene content and expression in their endosymbionts. Following assembly and comparative genome and transcriptome analyses, we discovered that these endosymbionts differed with respect to possible niche specific traits such carbon metabolism. Our analyses supported that genes related to carbon metabolism were acquired by horizontal gene transfer (HGT) from donor taxa which are present in termite’s hindgut community. In addition, our analyses supported that these endosymbionts have retained and expressed genes related to natural transformation (competence) and recombination. Taken together, the presence of genes acquired by HGT and a putative competence pathway supported that these endosymbionts are not cut-off from gene flow and that competence may be a mechanism by which members of the *Endomicrobium* can acquire new traits.

**Importance:** The composition and structure of wood, which contains cellulose, hemicellulose and lignin, prevents most organisms from using this common food source. Termites are a rare exception among animals, and they rely on a complex microbiome housed in their hindguts to use wood as a source of food. The lower termite *R. flavipes* houses a variety of protist and prokaryotes that are the key players the disassembly of lignocellulose. In this paper we describe the genomes and the gene expression profiles of five *Endomicrobium* endosymbionts living inside three different protist species from *R. flavipes*. Data from these genomes suggest that these *Endomicrobium* species have different mechanisms for using both carbon and nitrogen. In addition, they harbor genes that may be used to import free DNA from their environment. This process of DNA-uptake may cntribute to the high levels of horizontal gene transfer often seen in the *Endomicrobium* species.

## Introduction

Among the wood-feeding lower termites, symbiotic protists which reside in the hindgut are often colonized by endosymbionts (1–4). In *Reticulitermes* spp. termites both Oxymonadida (order) and Parabasilia (class) protists associate with endosymbiotic bacteria belonging to the genus *Endomicrobium* (phylum Elusimicrobia, class Endomicrobia) (2, 5–7). Members of *Endomicrobium* have been shown to comprise a significant portion of the core bacterial community in wood-feeding termites such as *R. flavipes* (8, 9). These endosymbiotic lineages are thought to have initiated their associations with hindgut protists approximately 70 - 40 million years ago (10) and arose from free-living relatives during multiple independent acquisition events (11). Vertical passage from one protist cell to its progeny, has resulted in co-speciation as inferred from congruent ribosomal RNA (rRNA) phylogenies (7, 10, 12, 13).

In addition to colonizing the cytoplasm of certain hindgut protist species, *Endomicrobium* spp. are ectosymbionts of protists (14) can be free-living as well (15)(11, 16, 17). Because of their distribution across these different niches, they provide an opportunity for studying bacterial genome evolution across different association lifestyles: free-living, endosymbiotic, and ectosymbiotic.

To determine differences two *Endomicrobium* species that are closely related but with distinct lifestyles, a previous study compared genomes of a free-living *Endomicrobium*, *E. proavitum* strain Rsa215 (16) isolated from *R. santonensis* (*R. flavipes*), and ‘*Candidatus* Endomicrobium trichonymphae’ strain Rs-D17 (3), an endosymbiont isolated from the cytoplasm of a *Trichonympha* from the termite *R. speratus* (3, 18). The findings suggested that the transition from the free-living state to an intracellular lifestyle involved genome reduction, similar to that of endosymbionts of sap-feeding insects and many obligate intracellular pathogens. However, the intracellular strain Rs-D17 also incorporated genes, possibly from other termite gut inhabitants, by horizontal gene transfer (HGT) (18). For example, the genome of ‘*Ca*. E.trichonymphae’ Rs-D17 appeared to have acquired several pathways including those that encode sugar and amino acid transporters and genes involved in amino acids biosynthesis (18). These findings suggested that, unlike the endosymbionts of sap-feeding insects, *Endomicrobium* species may not be completely cut-off from gene flow (18).

We expand upon these studies by presenting and comparing near-complete draft genomes and transcriptomes of three different *Endomicrobium* organisms, that were assembled from single protists cells of three different species that inhabit the hindgut of *R. flavipes*. One of these protists species, *Pyrsonympha vertens*, lives attached to the oxic gut wall (19, 20), while the other two *Trichonympha agilis* and *Dinenympha* species II, are found in the more anoxic hindgut lumen. In addition, *P. vertens* and *D.* species II are both Oxymonads while *T. agilis* is a Parabasalid.

The analyses indicate that these *Endomicrobium* have differences in gene content and expression, related to carbon usage and metabolism. And as seen previously in ‘*Ca*. E. trichonymphae’ Rs-D17, they have likely acquired genes from putative donor taxa that are commonly associated with termites. In addition, we describe data suggesting that these *Endomicrobium* have retained competence genes which may allow them to import exogenous DNA and that perhaps have contributed to HGT. Genes involved in this pathway are conserved across several *Endomicrobium* species and were expressed in the endosymbionts examined in this study.

## Methods

### Termite collection and species identification

*. R. flavipes* termites were collected using cardboard traps placed under logs for 2 to 4 weeks at the UConn Campus at Storrs, Connecticut (Longitude −72.262216, Latitude 41.806543) and their identity was verified as previously described (7) by amplifying and sequencing the mitochondrial cytochrome oxidase II gene. Termites were maintained in the lab with moistened sand and spruce wood that were initially sterilized.

### Single protist cell isolation

Termites from the worker caste were brought into an anaerobic chamber and their hindguts were dissected with sterile forceps. Hindguts were ruptured in ice-cold Trager’s Solution U (TU) (21) and washed three times by centrifuging in 500 ul of TU at 3,000 rpm in an Eppendorf microcentrifuge for 90 seconds. This washed cell suspension was then diluted 10-fold in TU buffer on ice. A 1 µl aliquot of the washed and diluted cell suspension was added to a 9 µl droplet on a glass slide treated with RNase AWAY® Reagent (Life Technologies) and UV light. Individual protist cells were isolated using a micromanipulator (Eppendorf CellTram® Vario) equipped with a hand-drawn glass capillary. Individual cells were washed three times in 10 µl droplets of TU via micromanipulation, transferring approximately 0.1 µl each time, and finally placed in 10µl molecular grade phosphate buffered saline (PBS), flash frozen on dry ice, and immediately stored at −80^°^C. Meta-data regarding these protist cell samples can be found as Supplementary Table 1 in S1 File.

### Whole genome and transcriptome amplification and sequencing

The metagenome (DNA) and metatranscriptome (cDNA) from individual protist cells and their associated bacteria were simultaneously amplified 12-24 hours after isolation. Cell lysis and amplification was performed using the Repli-g WGA/WTA kit (Qiagen). Cells were lysed using a Qiagen lysis buffer followed immediately by incubation on ice. Two samples from each lysed cell were taken and used in for whole genome amplification and whole transcriptome amplification. These were carried using the manufacturer’s standard protocol with exception that random hexamer primers were used to amplify DNA and cDNA. DNA and cDNA were sheared using a Covaris M220 ultra-sonicator™ according to the manufacturer’s protocol. WGA samples were sheared to a 550 bp insert size using 200 ng of DNA. WTA samples were sheared to a 350 bp insert size using 100 ng of cDNA. Sequencing libraries were prepared using the TruSeq Nano DNA Library Prep kit from Illumina^®^ according to the manufacturer’s protocol. Each sample was prepared with a forward and reverse barcode such that samples could be multiplexed on the same sequencing run. The samples were sequenced using an Illumina^®^ NextSeq 1×150 mid-output run and two NextSeq 1×150 high-output runs. Meta-data regarding amplicon yields can be found in Supplementary Table 1 in S1 File.

### Genomic read processing and assembly

Reads were preprocessed before assembly using BBmap (22). Reads were filtered for contaminating sequences by mapping reads to reference genomes of potential contamination sources such as human DNA, human associated microbiota, and organisms commonly used in our research laboratories. A list of references genomes used for contamination filtering is provided in Supplementary Table 2 in S1 File. Using BBmap scripts, adaptor sequences were trimmed from reads and last base pair of 151 bp reads was removed. Reads were then trimmed at both ends using a quality score cutoff of Q15. Homopolymers were removed by setting an entropy cutoff of 0.2, a max G+C cutoff of 90%, and by removing reads which possessed stretches of G’s equal to or greater than 23 bases long. In addition, reads which were below a minimum average quality of Q15 and/or 50 bases long were removed. Genomic reads were then normalized to a minimum coverage of 2X and a maximum coverage of 50X and then deduplicated using BBnorm. Genomic reads were assembled using the A5 assembly pipeline (23) on the KBase web server (24). Meta-data regarding metagenome and metatranscriptome reads numbers can be found in Supplementary Table 3 in S1 File.

### Genomic binning, draft genome assessment and annotation

Metagenomic assemblies from single protist host cells and their bacterial symbionts were binned using either 4mer or 6mer frequencies with VizBin (25) and scaffolds at least 1Kb in size. Clustered scaffolds in genomic bins of interest (low GC content) were selected in VizBin. Each scaffold from these bins were used in a blastn (26) search against previously sequenced Elusimicrobia genomes (Supplementary Table 4 in S1 File). Scaffolds which had a positive hit to other Elusimicrobia (at least 70% identity over a 1kb alignment) were retained in the draft genomes and scaffolds which did not have a significant hit to other Elusimicrobia genomes were used in a second blastn search against the non-redundant (NR) database. Scaffolds which had positive hits to other Elusimicrobia in the NR database were retained in the draft genomes. Draft genomes were iteratively polished with the program Pilon (27). These draft genomes were then assessed for contamination and completeness using CheckM which uses lineage specific marker genes to perform analyses (28). The resulting near-complete draft genomes were then annotated on the RAST Server using a customized RASTtk workflow with options selected to call insertion sequences and prophages (29, 30). Metabolic pathways pertaining to carbon metabolism, amino acid biosynthesis, vitamin biosynthesis, and peptidoglycan biosynthesis were reconstructed from the annotated genomes using pathways in the Kyoto Encyclopedia of Genes and Genomes (KEGG) (31).

### Analysis of ribosomal gene phylogeny and average nucleotide identities

Ribosomal 16S genes from each of the *Endomicrobium* spp. draft genomes were trimmed and aligned to references using MUSCLE (32), evolutionary models were tested and a Maximum likelihood (ML) phylogenetic tree was made using IQ-TREE (33). JSpeciesWS (34) was used for determining the genomic average nucleotide identities based on BLAST+ searches (ANIb) between the *Endomicrobium* spp. draft genomes and the genome of ‘*Ca*. Endomicrobium trichonymphae’ Rs-D17, which is a close relative (3).

Assembled 18S rRNA genes were retrieved from metagenome assemblies by performing a BLAST+ search using previously published 18S rRNA reference sequences for each protist species as queries (7). When possible, protist 18S rRNA genes were amplified using leftover DNA from WGA samples using universal primers 18SFU; 5’-ATGCTTGTCTCAAAGGRYTAAGCCATGC-3’ and 18SRU; 5’-CWGGTTCACCWACGGAAACCTTGTTACG-3’ (35) as previously described (7) and sequenced by Sanger sequencing. This confirmation PCR was done on samples TA21, TA26, and DS12. Assembled 18S rRNA genes were aligned to references using MUSCLE and a Maximum likelihood (ML) phylogenetic tree was generated using IQ-TREE with model testing.

### Detection of horizontally acquired genes

Genes that may have been acquired by horizontal gene transfer were identified using phylogenetic methods. Initially, protein sequences of genes of interest that were not shared across our draft genomes were aligned to references that spanned eight different bacterial phyla (including Acidobacteria, Bacteroidetes, Nitrospiraceae, Spirochaetes, Firmicutes, Actinobacteria, Proteobacteria, and group PVC) using MUSCLE and phylogenetic trees were generated using IQ-TREE with model testing. Gene trees were then compared to the 16S rRNA gene tree phylogeny (Supplemental Figure 3) to determine evolutionary incongruence.

### Analysis of genes involved in competence and recombination

Genes known to be involved in DNA uptake, competence, and recombination were identified in each *Endomicrobium* spp. draft genome based on their RAST annotations and homology to reference sequences. The distribution of these genes was then compared across draft genomes and references which included free-living relatives and other endosymbionts. To asses if these genes were complete and if the encoded proteins likely retained their putative functions, homologs of each gene were obtained from genomes of bacteria belonging to the phylum Elusimicrobia, aligned with MUSCLE, and phylogenetic trees were generated using IQ-TREE (33) with model testing, and support values generated using the “–abayes” and “–bb 1000” commands. The resulting phylogenetic trees were used along with the MUSCLE alignments to perform a dN/dS analysis using the program Codeml which is a part of the PAML and PAMLX packages (36, 37).

### Mapping transcriptome reads to draft genomes

RNA-seq metatranscriptome reads were quality trimmed and filtered as described above and error corrected in Geneious R11 (38) using BBNorm with default settings. To remove rRNA reads before mapping, rRNA sequences were identified from each metagenome assembly using RNAmmer (39) and reads were mapped to these, as well as, rRNA references from refseq (40), SILVA (41), and DictDb (42) databases using BBmap (22). Remaining metatranscriptome reads were then mapped to their respective *Endomicrobium* spp. draft genome in Geneious R11 using Bowtie 2 (43) with alignment type set to “End to End” and using the “Medium Sensitivity” preset. Expression levels were then calculated in Geneious R11, ambiguously mapped reads were excluded from the calculations. RPKM values for genes in each genome are given in Supplementary Tables 5 and 6 in S1 File

### Verification of *comEC* expression by RT-PCR

Primers were designed to amplify *comEC* from ‘*Ca*. Endomicrobium agilae’ in Geneious R11 using Primer3 (44) (Primers: endo_comec_F: 5’-ATTTGCCTGTGTTTGAGAGT-3’ and endo_comec_R: 5’-CCTGTTCCTGTGCTTTCAG-3’). Twenty termites were used to prepare RNA and cDNA samples for RT-PCR analysis. Termite hindguts were dissected and ruptured in TU on ice in an anaerobic chamber. Hindgut contents were washed with ice-cold TU three times at 3,000 rpm in an Eppendorf microcentrifuge for 90 seconds and then lysed in 1mL of TRIzol™ Reagent (Thermo Fisher Scientific). RNA was isolated per the manufacturer’s protocol and treated with TURBO™ DNase (Thermo Fisher Scientific) following the manufacturer’s protocol for 50µl reactions using ~5.6 mg of total RNA in a 20 ul volume. 20 ul of the DNase-treated RNA was then used as template for cDNA synthesis using SuperScript IV Reverse Transcriptase (Thermo Fisher Scientific) following the manufacturer’s protocol for first strand synthesis primed with random hexamers. The resulting cDNA was treated with *E. coli* RNaseH for 20 minutes at 37^°^C.

RT-PCR reactions were performed using the *Endomicrobium comEC* primers with RNaseH-treated cDNA serving as template and the no-RT control consisting of DNase-treated RNA that did not undergo cDNA synthesis. RT-RCR was performed using Phusion Polymerase (Thermo Fisher Scientific) with HF buffer and DMSO. Cycling conditions were: initial denaturation at 94^°^C for 3 minutes, followed by 35 cycles of 94^°^C for 45 seconds, annealing at 59^°^C for 30 seconds, and extension at 72^°^C for 45 seconds. Final extension was done 72^°^C for 10 minutes. Hindgut DNA (washed protists cell fractions from five hindguts in molecular grade Tris EDTA buffer) was used as a positive PCR control. RT-PCR products were visualized using a 1% agarose gel with ethidium bromide. Products were purified using the Monarch DNA gel purification kit (New England Biolabs), and Sanger sequenced.

### Data availability

Raw reads and assemblies will be submitted to NCBI GenBank under BioProject PRJNA644342.

## Results

### Phylogeny of protist hosts

Protist 18S rRNA genes were retrieved from metagenome assemblies and confirmed (when possible) independently by PCR and Sanger sequencing. A Maximum likelihood (ML) phylogenetic tree was made that indicated species of the protist cells used in this study were *Trichonympha agilis* (cells TA21 and TA26), *Pyrsonympha vertens* (cells PV1 and PV7), and *Dinenympha* species II (cell DS12) (Figure 1). These protist species have been previously confirmed to live associated with *R. flavipes*, the termite species used in this study.

**Figure 1.**
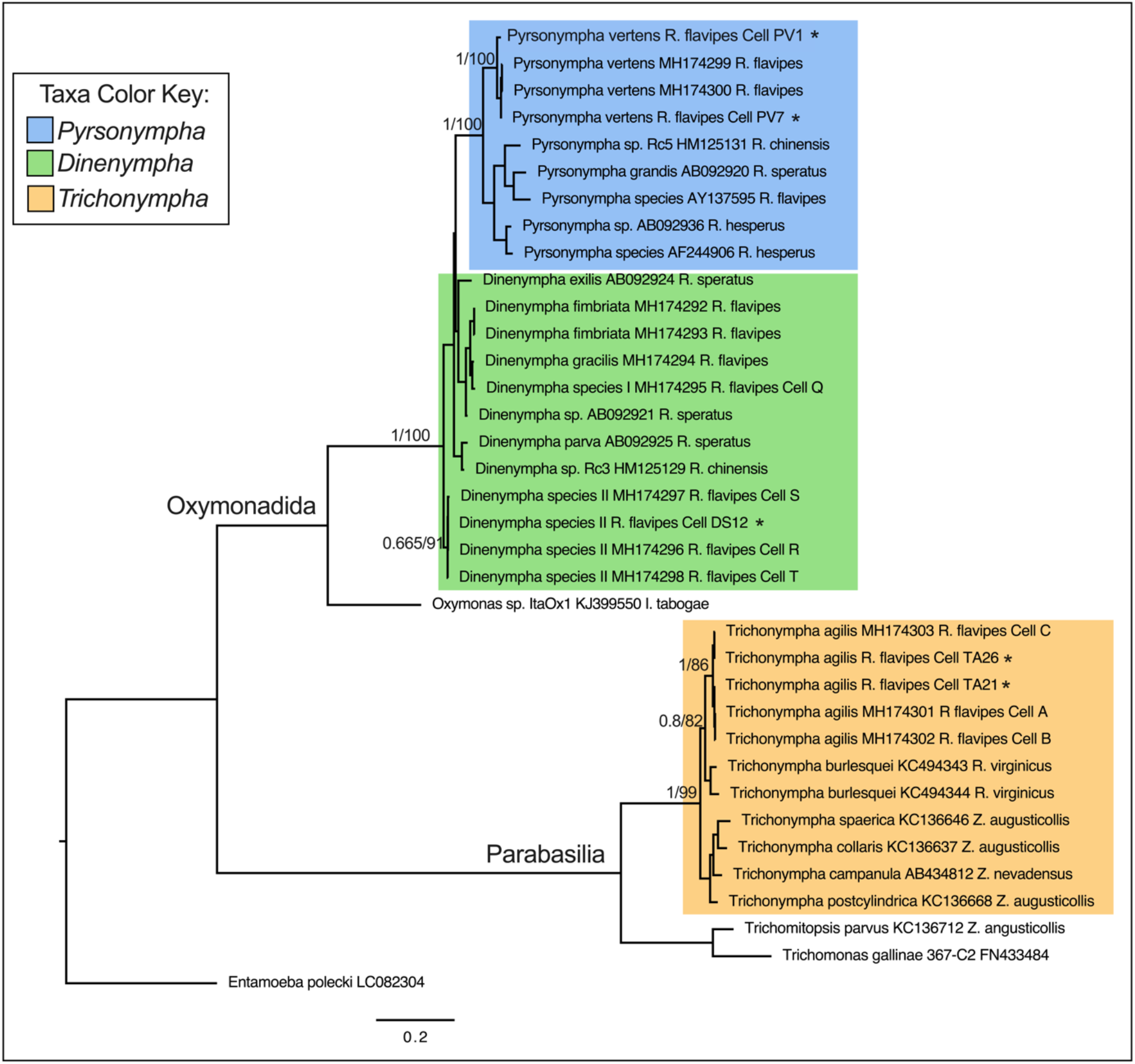
Protist 18S rRNA gene phylogeny. 18S rRNA genes were retrieved from single protist cell metagenome assemblies, aligned to references, and a Maximum likelihood (ML) phylogenetic tree was made using IQ-Tree using substitution model TIM2+G4. All 18S rRNA gene sequences obtained in this study (denoted by *) are shown grouped with their respective references. Branch support values represent the Bayesian posterior probability and Bootstrap support values respectively.

### *Endomicrobium* genome statistics

Five near-complete *Endomicrobium* genomes were obtained from single protist cell metagenomic assemblies. The five genomes ranged from 1.12 - 1.37 Mb in size, 35.3 – 36.6 % G+C, and 93.3 - 96.6% completeness (Figure 2A). To determine if these genomes were from the same or different *Endomicrobium* species, we calculated pairwise genomic s using an ANI score of 95% or greater as a marker for species-level cutoff (Figure 2B) (45).

**Figure 2.**
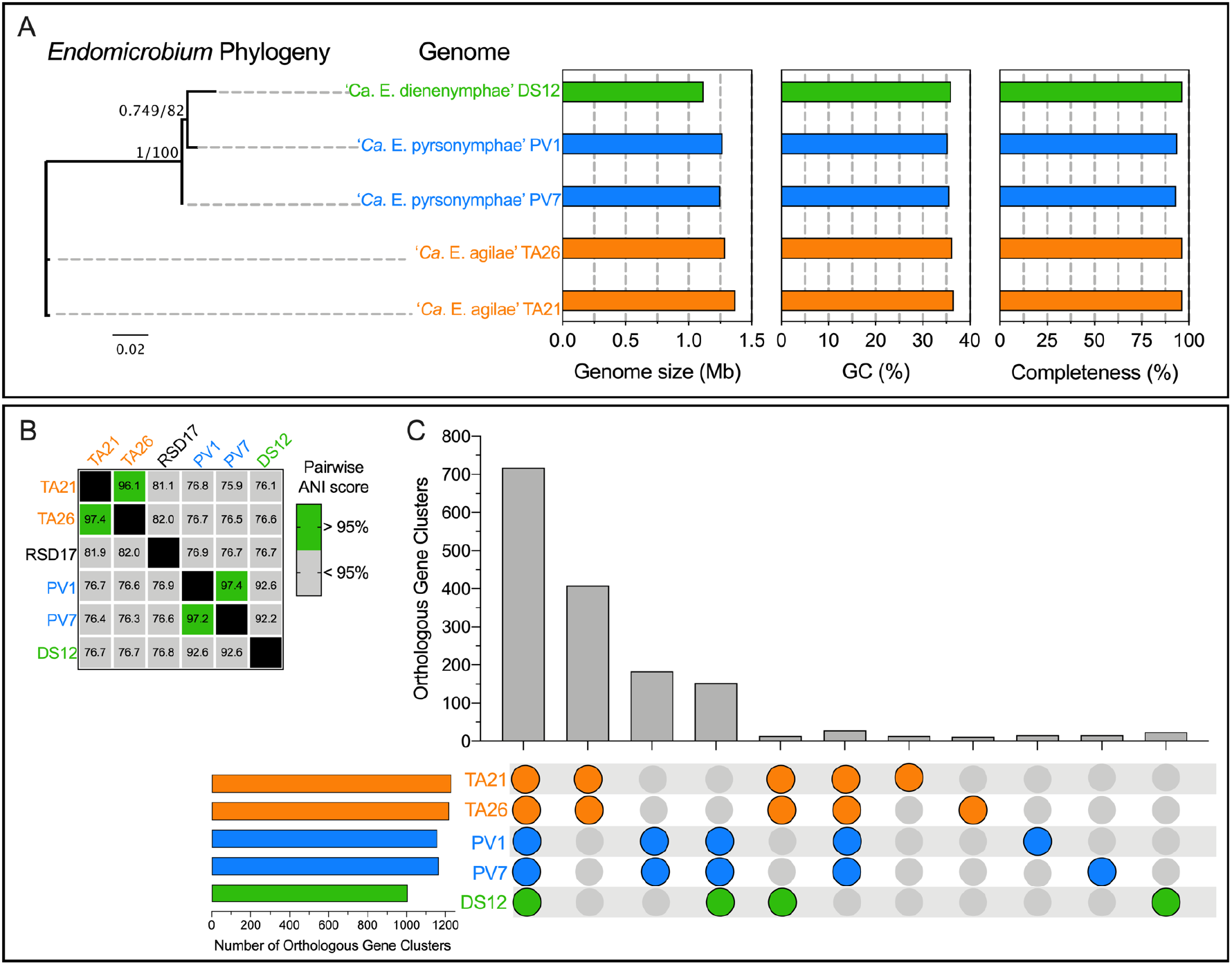
*Endomicrobium* draft genomes statistics, speciation, and shared gene content. (A) 16S rRNA gene Maximum likelihood tree (unrooted) of the three *Endomicrobium* species, genome sizes, percent G+C content, and estimated percent genome completeness. (B) Pairwise genomic ANI scores of *Endomicrobium* genomes obtained by this study and a previously sequenced relative Rs-D17. (C) UpSet graph of the number of orthologous gene clusters (OGCs) of protein coding sequences within and across each of the *Endomicrobium* draft genomes.

From *T. agilis* samples, we assembled two Endomicrobium draft genomes which had and ANI score of greater than 96% to one another but less than 90% to ‘*Ca*. E. trichonymphae’ Rs-D17 indicating that they are likely different species. Based on this analysis, we refer to the draft genomes as coming from ‘*Candidatus* Endomicrobium agilae’ TA21 and ‘*Candidatus* Endomicrobium agilae’ TA26. We also assembled two *Endomicrobium* genomes from *P. vertens* samples which had an ANI score greater than 97% identity to each another (Figure 2B) and whose 16S rRNA genes were greater than 98% identical to a previously described species, ‘*Candidatus* Endomicrobium pyrsonymphae’ (6), which is the Candidatus species designation that we use for PV1 and PV7. One additional *Endomicrobium* genome was assembled from *Dinenympha* species II. This genome did not share an ANI score greater than 95% to other *Endomicrobium* genomes and was thus given a new Candidatus species designation ‘*Candidatus* Endomicrobium dinenymphae’ DS12 (Figure 2B).

Individually, these *Endomicrobium* genomes contained between 1005 – 1230 orthologous gene clusters (OGCs), of which 717 were found in all five genomes (Figure 2C). Additionally, 409 OCGs were unique to ‘*Ca*. E. agilae’ TA21 and TA26 and another 183 OGCs were unique to ‘*Ca*. E. pyrsonymphae’ PV1 and PV7 (Figure 2C). Although the genome of ‘*Ca*. E. dinenymphae’ DS12 only had 24 unique OGCs, it shared 153 with ‘*Ca*. E. pyrsonymphae’ PV1 and PV7 (Figure 2C) which may reflect similar selective pressures for gene retention in their Oxymonad hosts *Dinenympha* and *Pyrsonympha*, or their more recent shared history, compared with the *Endomicrobium* (TA21 and TA26) that associated with the Parabasalid *T. agilis*.

### Biosynthesis of amino acids, vitamins and peptidoglycan

The presence of genes for the various function discussed below were detected by tblastn using the queries listed in Supplementary File 1 (Tables 8 and 9). When genes were not detected in this manner, read-mapping using Geneious and read-mapping using Megan (63) were done as well to identify reads from genes that may not have assembled into contigs. In general, each of the five *Endomicrobium* genomes assembled in this study had similar gene content for processes involved in the biosynthesis of amino acids (Supplementary Figure 1A), vitamins (Supplementary Figure 1B), and peptidoglycan (Supplementary Figure 1D). Each genome possessed complete pathways for alanine (from cysteine) aspartate, arginine, glutamine, glutamate, glycine (from imported serine), isoleucine, leucine, valine, lysine, tyrosine, phenylalanine and tryptophan biosynthesis (Supplementary Figure 1A). Interestingly, the *Endomicrobium* symbionts of Oxymonad protists (PV1, PV7 and DS12) lacked at least one gene in the biosynthesis pathway for histidine (*hisG*) (Supplementary Figures 1A and 2). The histidine biosynthetic pathway was complete in the genomes of ‘*Ca.* E. agilae’ TA21 and TA26 (Supplementary Figure 1A). Conversely, it is likely that the *Endomicrobium* symbionts of Oxymonad protists (PV1, PV7, and DS12) can make proline, while the symbionts represented by genomes TA21, TA26 and RsD17 cannot (Supplementary Figure 1A). The five genomes encoded incomplete pathways for the synthesis of cysteine and methionine. The three genomes isolated from Oxymonad protists encoded a methionine transporter (MetT) and all contained a gene encoding aB12-dependent methionine synthase system comprised of MetH and an activation protein MetH2 (Supplementary Figure 1A). Also incomplete in all five genomes were pathways for the synthesis of serine and asparagine. Each genome encoded a serine transporter SdaC, a proline transporter (ProT) and PV1, PV7, and DS12 each encoded a glutamate transporter, GltP (Supplementary Figure 1A).

The five *Endomicrobium* genomes also had similar gene content for processes involved in the biosynthesis of vitamins and co-factors, with the pathways to pantothenate, CoA, NAD and NADP being complete and other pathways being incomplete (Supplementary Figure 1B). Interestingly, the biotin biosynthesis pathways in the five genomes are missing just a single gene (*bioW*) needed to convert pimelate to pimelate-CoA suggesting that pimelate-CoA may be synthesized by another enzyme or imported (Supplementary Figure 1B). Several genes in the thiamine biosynthesis pathway were also missing in each of these genomes (Supplementary Figure 1B). As noted previously for *E. proavitum* and “*Candidatus* Endomicrobium trichonymphae” strain Rs-D17 the five genomes described here were also missing the steps in the folate pathway needed to make 4-aminobenzoate, which may be transported into the cells (18). The pathways for pyridoxine (B6) and vitamin B12 were also incomplete, though each of the five *Endomicrobium* genomes appeared to encode ABC transport systems for vitamin B12 and heme.

Regarding peptidoglycan synthesis, each *Endomicrobium* genome was missing an enzyme (BacA) that typically dephosphorylates undecaprenyl pyrophosphate. Since these different *Endomicrobium* species, including the free-living *E. proavitum*, are missing the same gene it may be that these bacteria utilize an alternate phosphatase to carry out the same function as BacA.

### Differences in Carbon Metabolism

Some of the more interesting differences between these *Endomicrobium* genomes pertained to their carbon metabolisms. Each of these five *Endomicrobium* genomes encoded relatively simple pathways for importing and using different wood-derived carbon sources. Each had a complete phosphotransferase system (PTS) for importing sugars. Present were two EIIA genes encoding sugar specific phosphorylation proteins most closely related to those of the mannose and fructose type EIIA proteins (Supplementary Figure 1C). Zheng et al. reported that *E. proavitum,* which contains a very similar PTS pathway, did not grow on mannose or fructose, but did grow on glucose, suggesting that glucose may be the carbohydrate transported by the PTS in that *Endomicrobium* species, and perhaps in the one described here as well (16, 18).

Based on the gene content in the five genomes analyzed here, carbon sources capable of being catabolized by endosymbiotic *Endomicrobium* species may often differ from each other and from their free-living relatives. For example, ‘*Ca.* E. agilae’ TA21 and TA26 encoded all the genes necessary to import and use both glucuronate and glucose-6-phosphate (Figure 3A & 3B). The closely-related ‘*Ca*. E. trichonymphae’ Rs-D17 (3) also contained these genes. Interestingly, genome analyses suggest that these two carbon sources cannot be used by the other *Endomicrobium* species studied here which lack the glucuronate transporter ExuT, the glucuronate isomerase UxaA and the glucose-6-phosphate transporter UhcP. The other *Endomicrobium* genomes encoded either arabinose (‘*Ca*. E. pyrsonymphae’ PV1 and PV7) or xylose (‘*Ca*. E. dinenymphae’ DS12) import and catabolism proteins that were not encoded in the TA21, TA26, *E. proavitum* or ‘*Ca*. E. trichonymphae’ Rs-D17 genomes. (Figures 4A & 4B; Figures 5A & 5B, respectively). Transcriptome data indicated that each of the genes involved in these carbon usage pathways were expressed in the respective *Endomicrobium* while they resided in their protist hosts (Figures 3C, 4C, & 5C). Metabolites from these carbon sources are typically fed into both the non-oxidative pentose phosphate pathway and glycolysis, both of which are complete in the five genomes described here (Figures 3B, 4B, 5B, & Supplementary Figure 1C).

**Figure 3.**
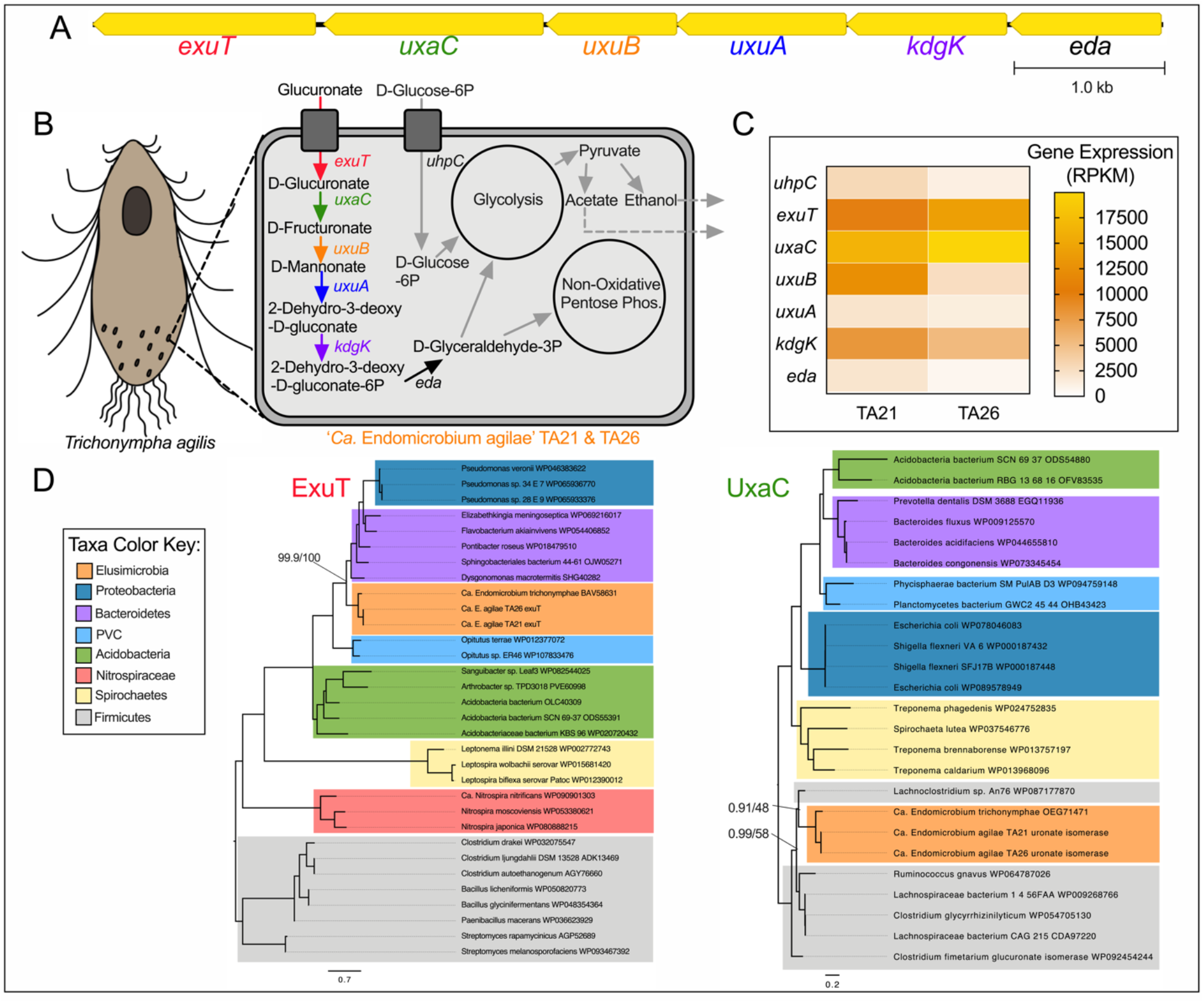
Carbon metabolism and HGT in ‘*Ca*. Endomicrobium agilae’. (A) Gene neighborhood of the genes involved in the metabolism of glucuronate in the ‘*Ca*. Endomicrobium agilae’ TA21 and TA26 genomes. (B) Diagram of a protist host and an *Endomicrobium* cell showing the inferred metabolic conversions of carbon sources based on gene content data. (C) Gene expression data of genes of interest (rows) pertaining to carbon metabolism in ‘*Ca*. Endomicrobium agilae’ TA21 and TA26 (columns). (D) Maximum likelihood phylogenetic trees of amino acid sequences of the transporter (ExuT, using substitution model LG+F+G4) and isomerase (UxaC, using substitution model LG+I+G4) in the glucuronate metabolism pathway. Support values represent the Bayesian posterior probability and Bootstrap support values respectively.

**Figure 4.**
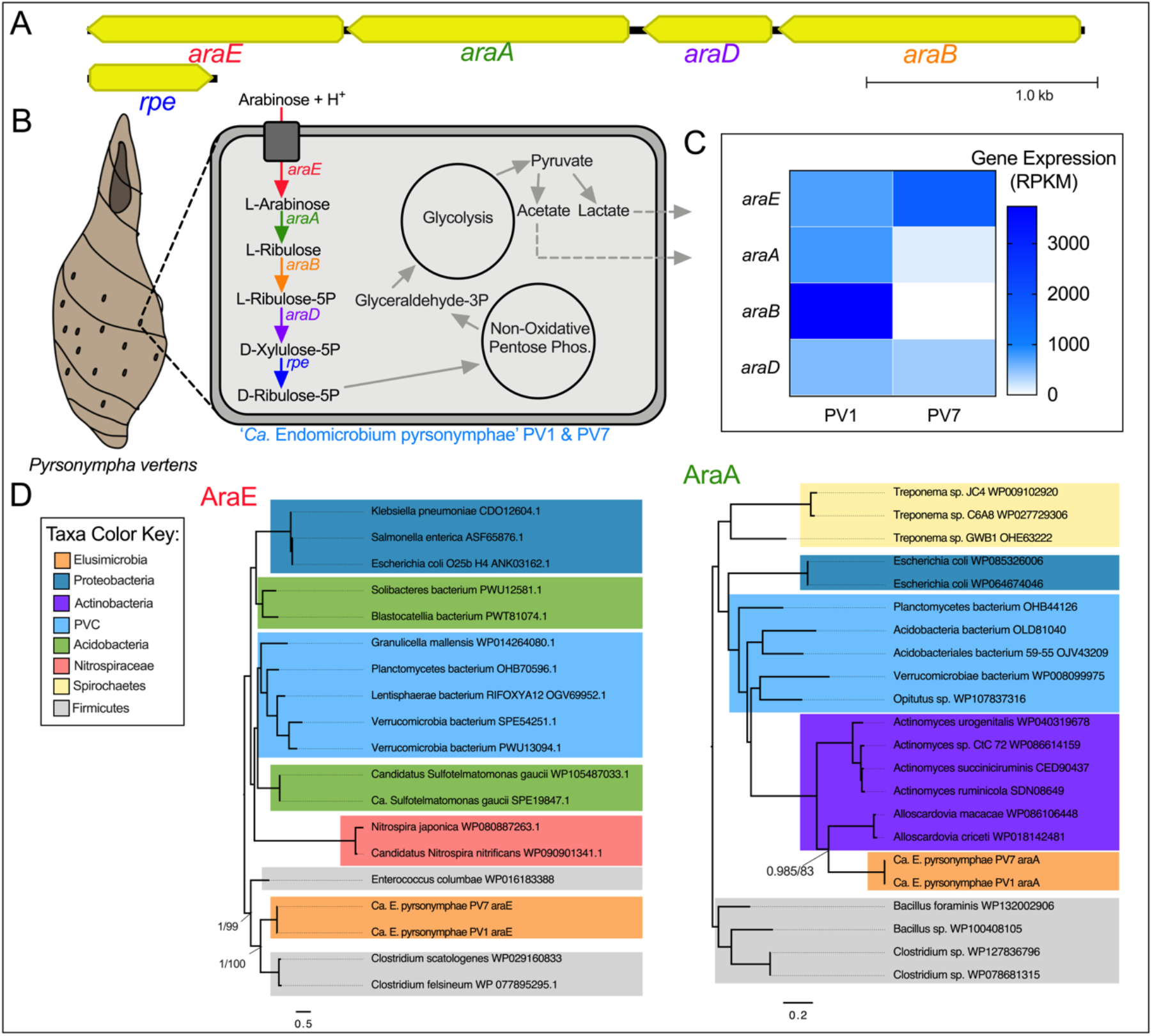
Carbon metabolism and HGT in ‘*Ca*. Endomicrobium pyrsonymphae’. (A) Gene neighborhood of the genes involved in the metabolism of arabinose in the ‘*Ca*. Endomicrobium pyrsonymphae’ PV1 and PV7 genomes. (B) Diagram of a protist host and an *Endomicrobium* cell showing the inferred metabolic conversions of carbon sources based on gene content data. (C) Gene expression data of genes of interest (rows) pertaining to carbon metabolism in ‘*Ca*. Endomicrobium pyrsonymphae’ PV1 and PV7 (columns). (D) Maximum likelihood phylogenetic trees of amino acid sequences from the transporter (AraE, using substitution model LG+F+G4) and isomerase (AraA, using substitution model LG+I+G4) in the arabinose metabolism pathway. Support values represent the Bayesian posterior probability and Bootstrap support values respectively.

**Figure 5.**
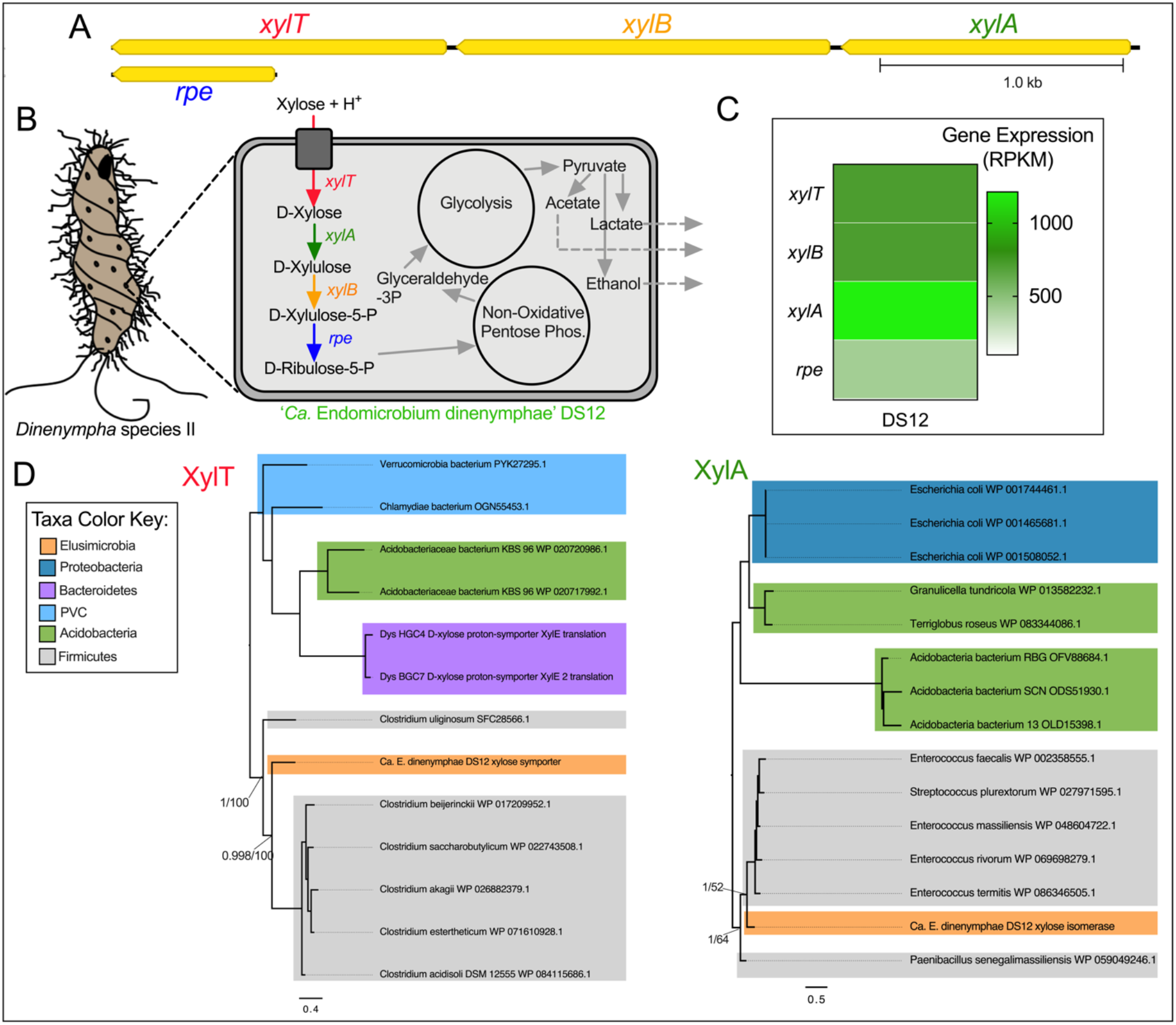
Carbon metabolism and HGT in ‘ *Ca*. Endomicrobium dinenymphae’. (A) Gene neighborhood of the genes involved in the metabolism of xylose in the ‘*Ca*. Endomicrobium dinenymphae’ DS12 genome. (B) Diagram of a protist host and an *Endomicrobium* cell showing the inferred metabolic conversions of carbon sources based on gene content data. (C) Gene expression data of genes of interest (rows) pertaining to carbon metabolism in ‘*Ca*. Endomicrobium dinenymphae’ DS12 (column). (D) Maximum likelihood phylogenetic trees of amino acid sequences from the transporter (XylT, using substitution model LG+F+G4) and isomerase (XylA, using substitution model LG+G4) in the xylose metabolism pathway. Support values represent the Bayesian posterior probability and Bootstrap support values respectively.

Other likely differences in carbon metabolism of these *Endomicrobium* species, included the production of fermentation end products (Supplementary Figure 1C). Analysis of the five *Endomicrobium* genomes suggested that following glycolysis, pyruvate can be fermented to acetate, however only the genomes of ‘*Ca.* E. agilae’ and ‘*Ca*. E. dinenymphae’, encoded AdhE which can convert acetate to ethanol. (Figures 3B, 4B, & 5B). In addition, genes encoding lactate dehyrodenase (LdH) were in the genomes of both ‘*Ca*. E. dinenymphae’, and ‘*Ca*. E. pyrsonymphae’, but not ‘*Ca.* E. agilae’ or ‘*Ca*. E. trichonymphae’ Rs-D17 (Figures 3B, 4B, & 5B). Differences in these fermentation pathways between free-living *E. proavitum* and ‘*Ca*. E. trichonymphae’ Rs-D17 were described earlier by Zheng et al. (18).

Previous studies identified genes acquired by horizontal gene transfer (HGT) in other *Endomicrobium* species (18), therefore, we tested whether HGT could, at least in part, explain the differences seen in carbon metabolism across the genomes presented in this study. Phylogenetic trees were made for each of the transport and isomerase proteins in the glucuronate, arabinose and xylose degradation pathways (Figures 3D, 4D, & 5D) and the phylogenies compared to the *Endomicrobium* 16S rRNA gene phylogeny to determine if they were congruent (Supplementary Figure 3). In each case, these phylogenies were not congruent, suggesting that these genes were acquired by HGT (Figures 3D, 4D, & 5D). Likely donor taxa include Bacteroidetes, Actinobacteria, and Firmicutes (Figures 3D, 4D, & 5D), which are all part of the hindgut community of *R. flavipes* (8). Similar data supporting HGT in ‘*Ca*. E. trichonymphae’ Rs-D17 have been reported and suggest that *Endomicrobium symbionts* are not cut off from gene flow and HGT (18). This is in contrast to the older endosymbionts of sap-feeding insects, which are traditionally thought to experience little to no gene flow, however recent analyses suggested that HGT may occur more frequently than previously thought in these symbionts (46).

### Natural transformation and competence as a possible mechanism for acquiring genes

Analyses of sequenced genomes of endosymbiotic *Endomicrobium* lineages indicate that acquisition of genes by HGT is relatively common. Thus, their genomes could reveal insights into the mechanisms by such genes were acquired. Interestingly, compared to other endosymbionts, the *Endomicrobium* genomes were enriched in genes related to the uptake of exogenous DNA and recombination (natural transformation/competence) (Supplementary Figures 4A and 4C). Of special interest are the *Endomicrobium* genes *comEC*, *comEB*, *comF*, *comM*, *ssb*, *drpA*, and *recA* which are all involved in natural transformation in bacteria such as *Vibrio cholerae* (47).

The dN/dS analyses of these genes supported the hypothesis that selection was acting to maintain the amino acid sequences of their corresponding gene products (dN/dS < 1.0) with the exception of *ssb* from TA21 (Figure 6A). In addition, transcriptome analysis indicated that these genes were expressed (Figure 6B). Expression of *comEC*, which encodes a transporter that imports single stranded DNA across the inner-membrane and into the cytoplasm of Gram-negative bacteria (47, 48), was verified by RT-PCR and sequencing of *Ca*. Endomicrobium agilae using *comEC* specific primers on a protist cell fraction sample prepared from 20 worker termite hindguts (Figure 6C). Together these data support the hypothesis that genes involved in this competence pathway are both conserved and expressed in these *Endomicrobium* symbionts of hindgut protists of *R. flavipes*.

**Figure 6.**
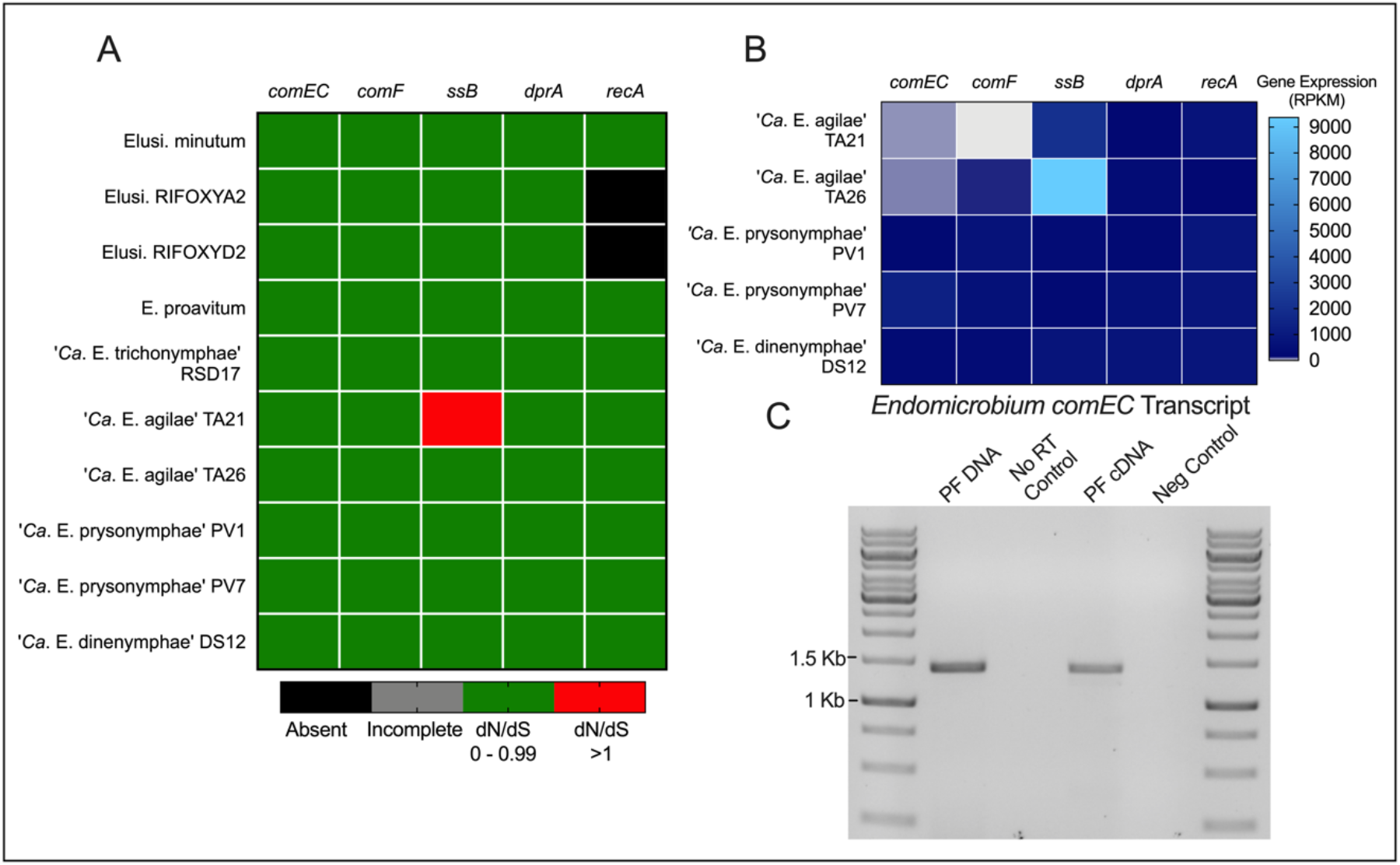
Analysis of genes involved in a putative competence pathway in *Endomicrobium* spp. (A) Heatmap showing the results of dN/dS analyses of genes involved in competence and recombination (columns) from *Endomicrobium* spp. and *Elusimicrobium* relatives (rows). (B) Gene expression data of those genes (columns) in the *Endomicrobium* spp. (rows) presented in this study. (C) RT-PCR gel image of *Endomicrobium comEC* transcript. Samples consisted of protist fraction (PF) DNA (positive control), No RT Control, PF cDNA, and molecular grade water (Negative control). Accession numbers for reference genomes used can be found in Supplementary Table 2.

The competence genes discussed above are involved in the translocation of single-stranded DNA across the inner membrane of Gram-negative bacteria and subsequent recombination. Also present in the genomes of all five *Endomicrobium* species analyzed in this study are genes which encode proteins that are similar to Type IV pilins. The TA21 and TA26 genomes contained a large chromosomal region devoted to Type IV Tad-like pilus synthesis as does E*. proavitum* and *E. minutum*. The bacterium *Ca*. E. trichonymphae’ Rs-D17 has a similar region, but it appears that many of the genes have become pseudogenes. The *P. vertens* and *Dinenympha* species II *Endomicrobium* symbionts had genes encoding pilins similar to the Type II PulG pilins. Some pilins from classes Type-IV and Type-II can bind and import double-stranded DNA across the outer membrane and have been shown to work in conjunction with ComEC-type proteins (47, 49). In addition, all five genomes possessed a pre-pilin peptidase (PilD). A graphical summary of these findings along with a model of how competence may work in these organisms is provided as Supplementary Figure 6. A list of genes and their putative function can be found in Supplementary Table 7 in S1 File.

### Transcriptome analysis of *Endomicrobium* populations inside single protist cells

Transcriptome analyses of the *Endomicrobium* populations inside single protist cells (from which the five genomes were derived) revealed similar gene expression profiles with a few notable exceptions (Figure 7). While our sample size for this work was necessarily small and while we were unable to do time-resolved sampling of the hindgut community from single termites, some general trends did appear in the transcriptomic data. Among COG categories, which are quite broad, the expression by each endosymbiont population was relatively similar with one exception being that there was higher expression of genes related to carbohydrate transport and metabolism in *Trichonympha* hosts (TA) compared to the Oxymonad hosts (PV and DS12) (Figure 7A).

**Figure 7.**
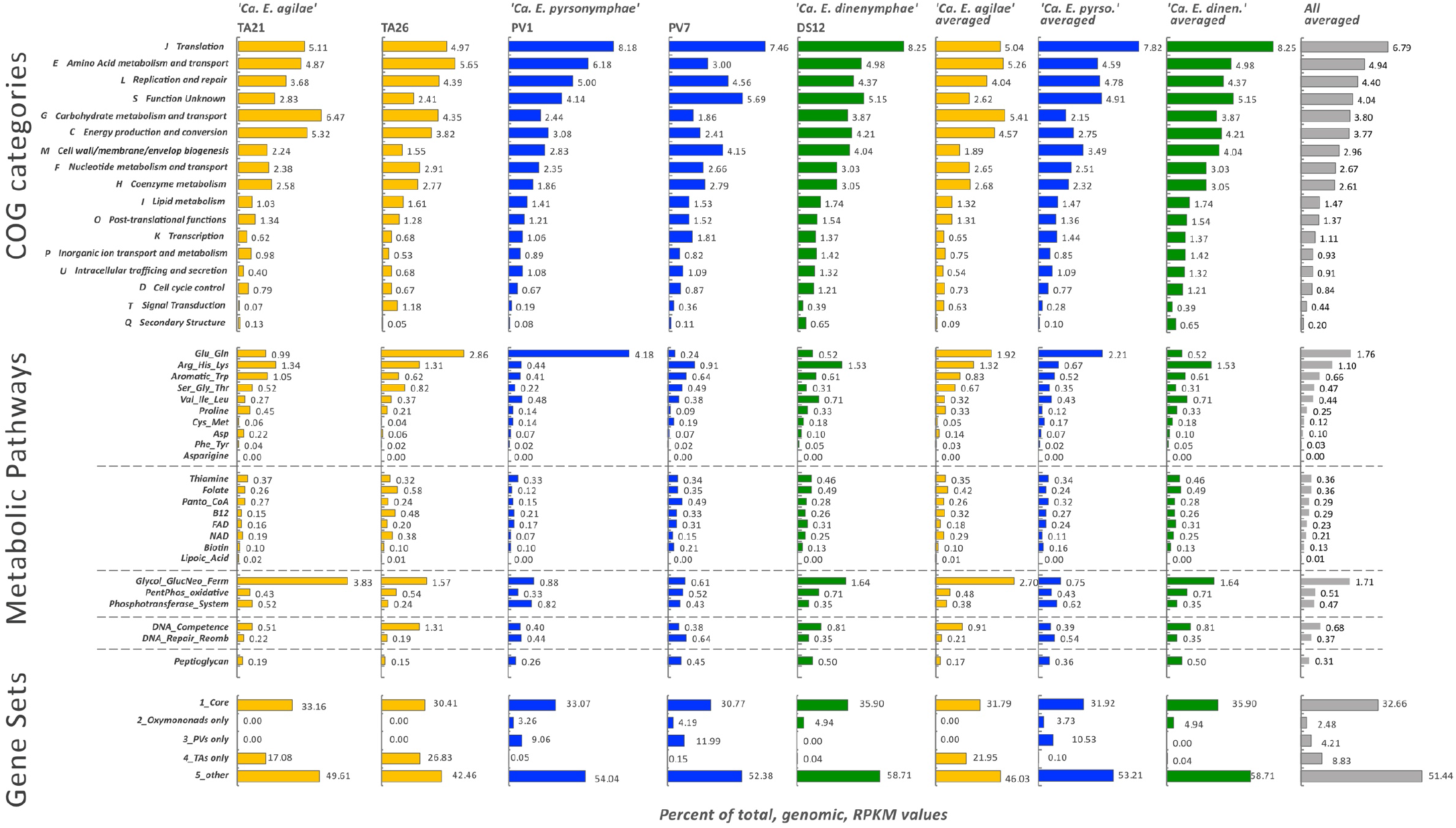
Transcriptome analysis of *Endomicrobium* populations from individual protist cells. (Top) Expression analysis of genes grouped into clusters of orthologous groups (COG) functional categories. (Middle) Expression analysis of genes in certain metabolic pathways pertaining to amino acid and cofactor biosynthesis, DNA processing, and peptidoglycan biosynthesis. (Bottom) Expression of genes grouped by their distributions among the different *Endomicrobium* species. RPKM values for individual cells, and averages of genera and all cells are in the various columns.

Analyses which focused on narrower categories such genes in related biosynthetic pathways, carbohydrate transport and break down, peptidoglycan synthesis and DNA uptake and repair revealed further differences not only between the endosymbionts of different protist species but even between the populations of endosymbionts of individual protist cells of the same type (Figure 7B). This is demonstrated by the differences related to the expression of genes in the glutamine and glutamate biosynthesis pathway (Figure 7B). Overall this pathway is more highly expressed by the endosymbionts of *Pyrsonympha* hosts compared to those in other protist species but there was variation in the expression of this pathway between individual protist host cells. For example, in host cell PV1 this pathway represented 4.2% of the total transcriptome reads, mostly from the gene *glnN* encoding a glutamine synthase, whereas in host cell PV7 it comprised only 0.2% (Figure 7B). Similar variation in this pathway was seen in the TA21 and TA26 transcriptomes. This data suggests that even populations of the bacterium residing in different host cells may not be expressing the same functions at any given point in time.

Core genes, which were shared between all five of the *Endomicrobium* species, represented an average of 30% – 36% of the transcription of each endosymbiont population (Figure 7C). The expression of genes that were specific to each *Endomicrobium* species ranged from 11% in ‘Ca. E. pyrsonymphae’ to 22% in ‘Ca. E. agilae’ indicating that there is likely differential gene content and gene expression endosymbionts of different protist host species (Figure 7C). Transcriptomic data are in Supplementary Tables 5 and 6 in S1 File

## Discussion

Single-cell protist metagenomics has enabled the assembly of genomes from several protist-associated bacterial symbionts from termites (1, 3, 4, 50, 51). In this study, we present near-complete draft genomes and transcriptomes of five endosymbiotic *Endomicrobium* samples, from three different protist species. These endosymbionts displayed differences with regards to their gene content and expression. For example, these organisms possessed different carbon usage pathways. One hypothesis that may explain such differences in carbon utilization is that different carbon sources may be provided to the *Endomicrobium* endosymbionts as by-products of the protist hosts’ hydrolysis and fermentation of the polysaccharides that originated in wood. For example, glucuronate may be present in the cytoplasm of *Trichonympha* spp. because they possessed the enzymes needed to cleave those monomers from polysaccharides found in wood, whereas the other protists, *P. vertens* and *D*. species II, may not be able to generate such monomers, or they may use them for other purposes (see below). If true, this suggests that there may be specialization among the protists with regards to polysaccharide hydrolysis in the hindgut of wood-feeding termites. A recent study demonstrated a division of labor among symbiotic protist species in a different termite, *Coptotermes formosanus*, where certain protist species produce different hydrolytic enzymes to degrade polysaccharides found in wood (52). These differences in protist functions may also explain why there was higher expression of genes related to carbohydrate transport and metabolism in the endosymbionts of *Trichonympha* hosts compared to the other protist species (Figure 7).

However, an alternative hypothesis is that metabolites can be partitioned within the host and some are specifically provided to certain symbionts. This, if true, may allow the host to control endosymbiont population densities through selective carbon source provision. Such host control of carbon provisioning is thought to operate in nitrogen-fixing root nodule symbioses ensuring that bacterial symbionts continue to provide fixed nitrogen in return for plant-provided carbon. In support of the second hypothesis, the membrane-embedded symbiont ‘*Ca*. Desulfovibrio trichonymphae’ which co-colonizes the same *Trichonympha* host as ‘*Ca*. E. trichonymphae’ Rs-D17, uses malate and citrate as carbon sources whereas its co-inhabitant likely uses glucuronate and glucose-6-phosphate (51).

Evidence suggests that *Endomicrobium* species have acquired genes by HGT and some of the donor taxa may include termite-associated bacteria. Endosymbiotic lineages of *Endomicrobium*, and their free-living relatives, possess many genes involved in DNA uptake, repair, and recombination (Supplementary Figure 4). Our analyses showed that the genes *comEC*, *comEB*, *comF*, *comM*, *ssb*, *drpA*, and *recA* are usually conserved within the Elusimicrobia phylum and were expressed in the endosymbiotic *Endomicrobium* species characterized in this study (Figure 6). Collectively these genes have been shown to be involved in the translocation of single stranded DNA across the inner membrane of other Gram-negative bacteria and in homologous recombination (47). The gene *comEC*, in particular, has an important function in this process as it encodes an essential part of the DNA transporter (47, 48). Using both transcriptome data, RT-PCR, and sequencing we were able to show that ‘*Ca*. Endomicrobium agilae’ *comEC* is expressed in these endosymbionts (Figure 6B and 6C). Collectively the genes involved in the competence pathway comprised between 0.4% to 1.3% of the total transcriptome reads of each endosymbiont population (Figure 7). These data suggest that *Endomicrobium* species may have the ability to become competent which may allow them to acquired DNA from the wider termite gut community and could result in HGT.

It is not clear how these organisms transport DNA across their outer membranes. None of these *Endomicrobium* species possessed all the components of a Type IV pili-based DNA-translocation system, but ‘*Ca*. E. agilae’ TA21 and TA26 contained a near-complete Type IV *tad* system which may allow DNA uptake (Supplemental Figures 4 and 5). It is also puzzling why each of the five genomes have retained *pilD,* a prepilin peptidase as well as genes encoding Type II and Type IV pilins. These may carry out some function in DNA-uptake or they may be non-functional and are in the process of being lost. If competence is a common trait among the *Endomicrobium*, this could explain why these organisms have many genes acquired through HGT and this capability may allow for rapid adaptation to new and diverse niches. Because hindgut protists phagocytize wood, wood-associated bacteria and perhaps free-living bacteria (53), this may be a route through which endosymbiotic *Endomicrobium* could be exposed to exogenous DNA.

However, it is worth noting that competence is not the only plausible avenue for DNA acquisition in these endosymbionts. HGT could also occur by bacteriophage transduction, conjugation, or other routes. Several lines of evidence indicate that these endosymbionts are susceptible to molecular parasites, such as bacteriophages and plasmids. Previous studies have reported that *Endomicrobium* species possessed several intact defense mechanisms to combat molecular parasites such as CRISPR-Cas and restriction-modification systems (54, 55). The *Endomicrobium* species sequenced in this study also contained those defense systems. The complete genome sequence of a bacteriophage of an endosymbiont (‘*Candidatus* Azobacteroides pseudotrichonymphae’) of a termite hindgut protist has previously been published, indicating that phage infection is not limited to *Endomicrobium* endosymbionts and may be common in termite hindguts (56).

Our analysis of *Endomicrobium* genomes and transcriptomes obtained from single protist cell metagenomes, highlighted several important differences across protist hosts which have led to hypotheses that warrant further investigation. In each case, the major hurdle of testing these is the current inability to culture the protists hosts which restricts their experimental tractability. However, the use of additional and new-omics approaches should further our understanding of these symbioses by focusing on the hosts’ and their symbionts’ genes, mRNA, metabolic and protein contents (57–62).

## Acknowledgments

This research was funded by the National Science Foundation (NSF) division of Emerging Frontiers in Research and Innovation in Multicellular and Inter-kingdom Signaling. Award number 1137249 (DJG and JG).

## Supplementary Figures and Legends

**Supplementary Figure 1.**
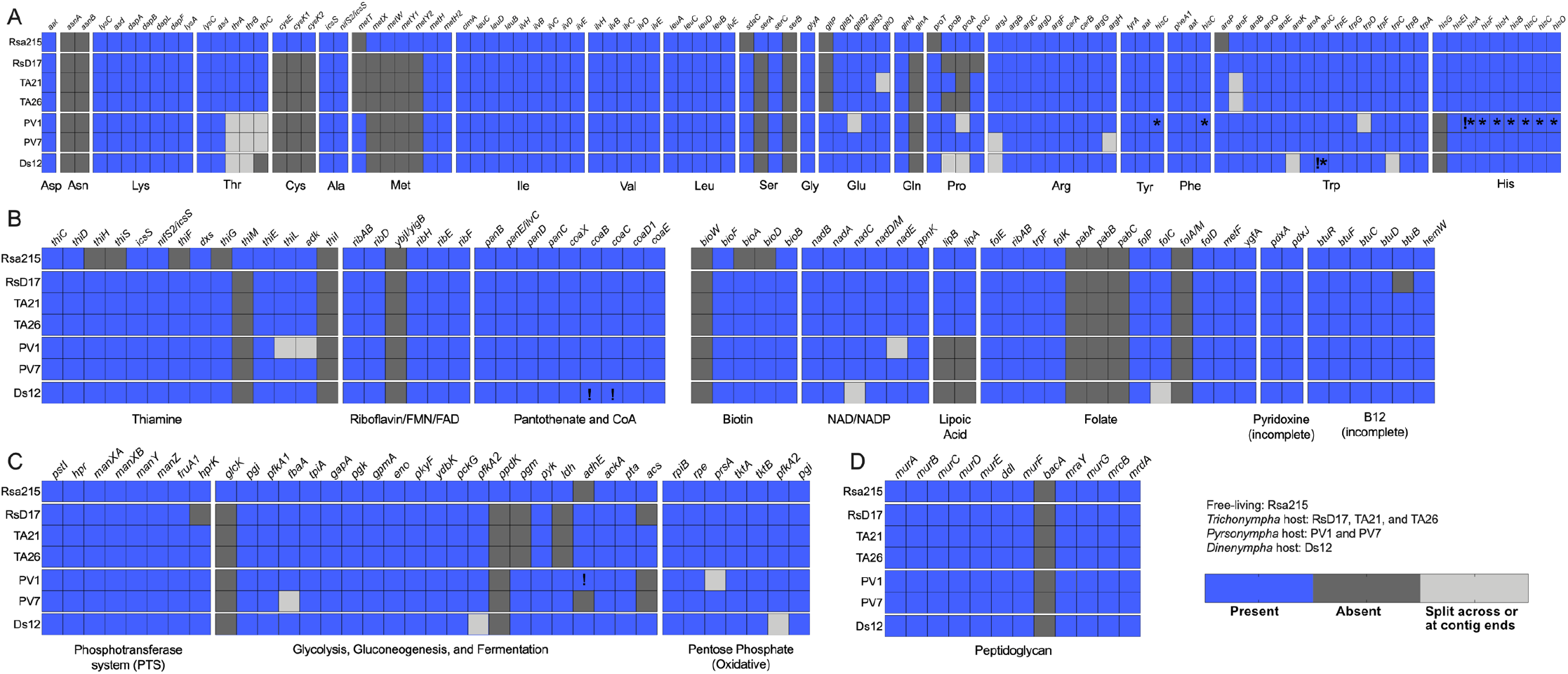
Gene content in *Endomicrobium* spp. genomes regarding metabolic functions and cell wall biosynthesis. (A) Gene content of amino acid biosynthesis pathways. (B) Gene content of vitamins and co-factor biosynthesis pathways. (C) Genes involved in central metabolism. (D) Gene content of peptidoglycan biosynthesis. Note that the genes marked with a “*” or a “!” were not in the final assemblies, but their reads were detected by either Megan (!) or by read mapping onto the same gene from another closely related organisms (*) or both (!*) (see Supplementary Figure 2 as an example).

**Supplementary Figure 2.**
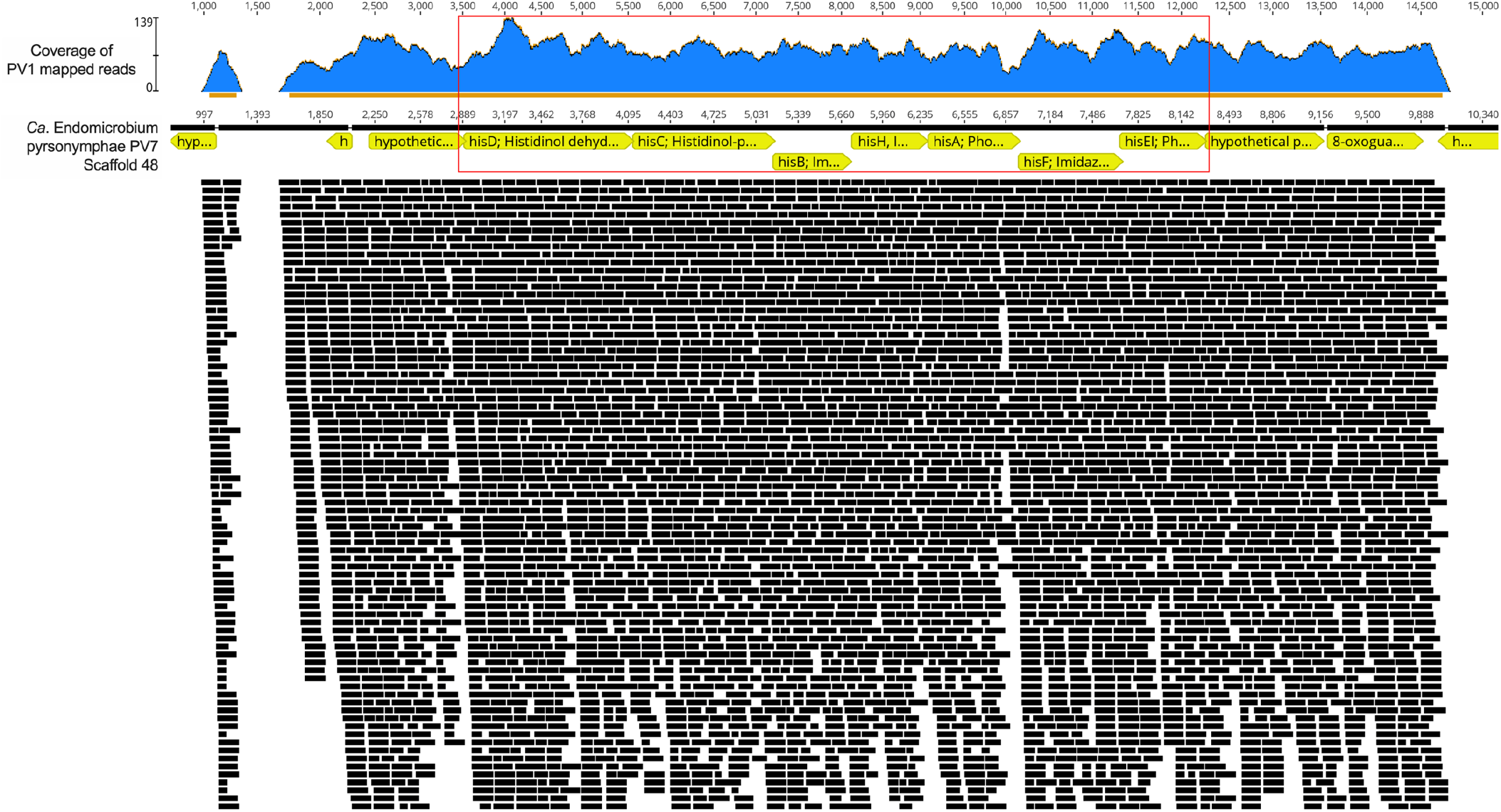
Mapping coverage of the histidine biosynthesis pathway in ‘*Ca*. Endomicrobium pyrsonymphae’ PV1. Metagenomic reads from sample PV1 were mapped to the draft genome of ‘*Ca*. Endomicrobium pyrsonymphae PV7’. The resulting coverage of these genes indicate that ‘*Ca*. Endomicrobium pyrsonymphae PV1’ encoded the histidine biosynthesis pathway and that the reason those genes are missing from the draft genome is likely due to an artifact of assembly or binning.

**Supplementary Figure 3.**
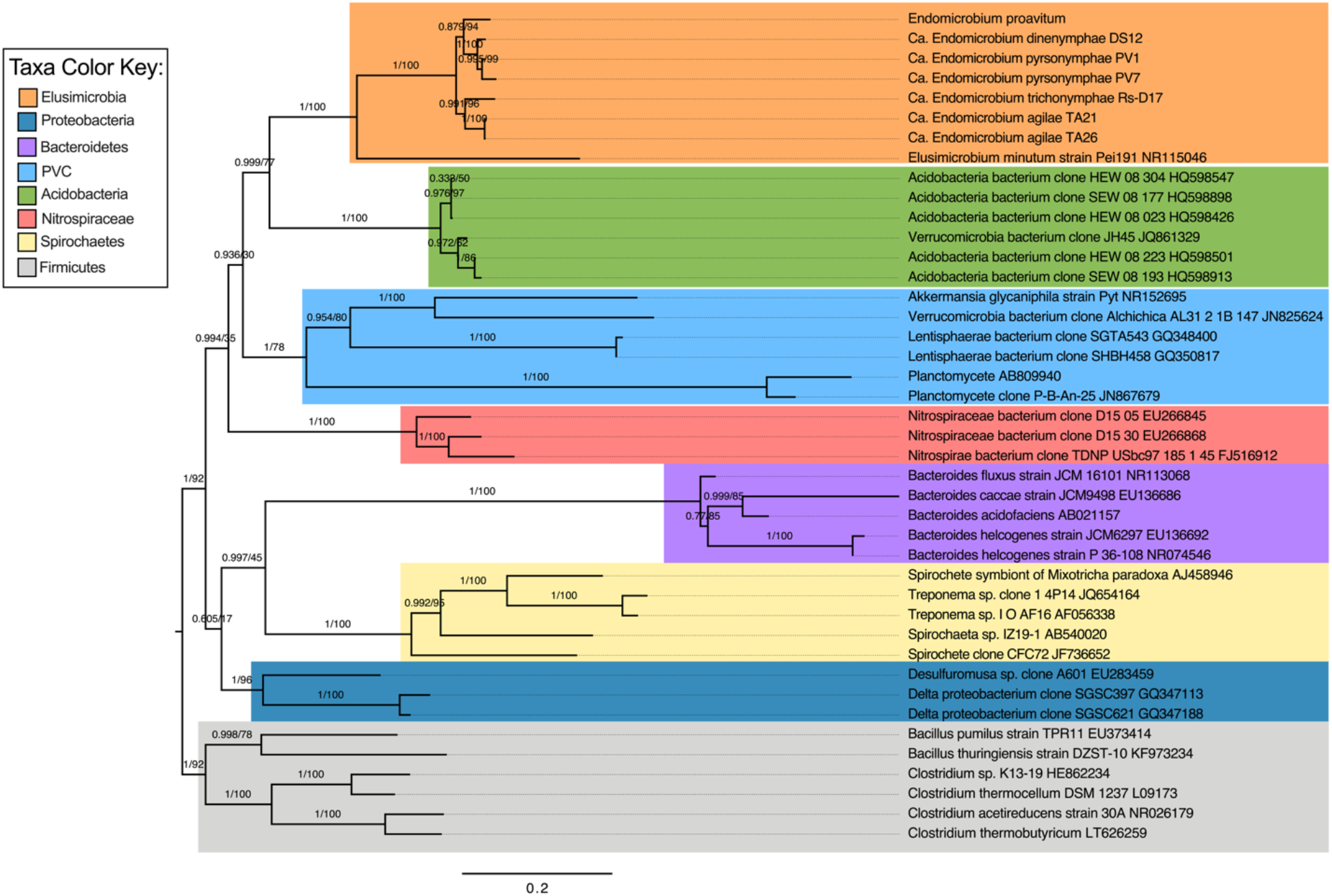
16S rRNA phylogeny of Elusimicrobia with respect to other phyla. Maximum likelihood phylogenetic tree of 16S rRNA genes (using substitution model TPM3u + G4) which was used a marker-gene to establish an organismal phylogeny of the Elusimicrobia phylum. This phylogeny was used to determine incongruence between the 16S rRNA gene and other genes of interest that may have been acquired via HGT by the *Endomicrobium* species.

**Supplementary Figure 4.**
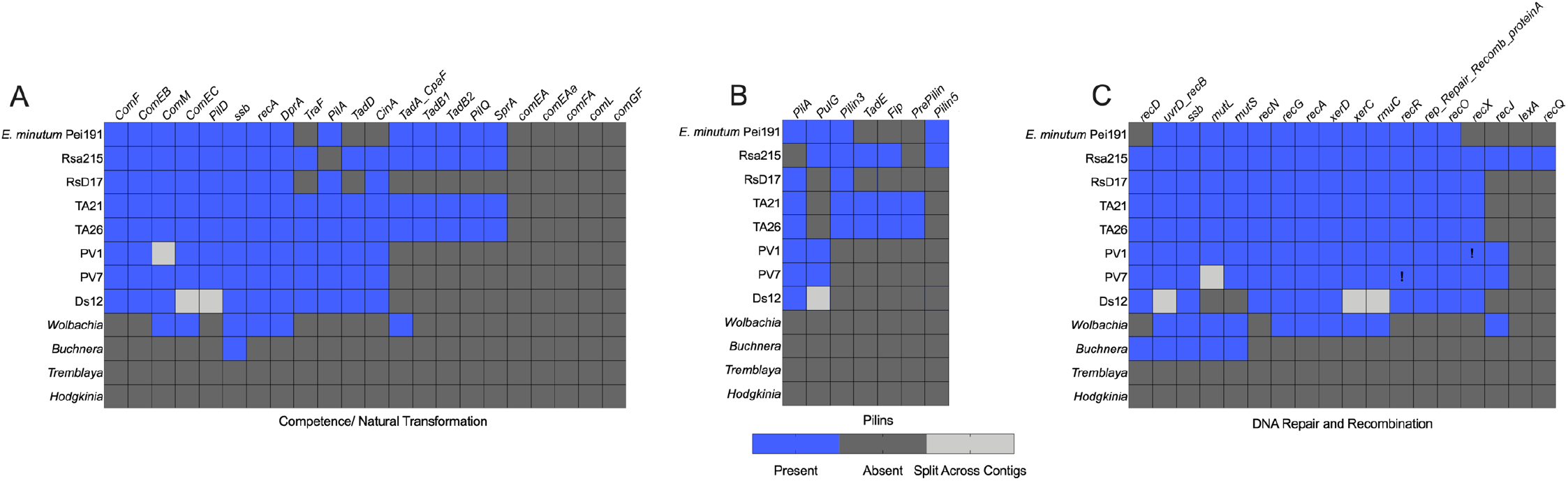
Gene content of regarding genes involved in natural transformation, pilus assembly, and DNA recombination/repair. (A) Presence and absence matrices of genes involved in natural transformation, (B) pilus assembly, and (C) DNA repair and recombination found the endosymbiotic *Endomicrobium* spp. genomes. The presence of these genes was investigated in their free-living relatives (Rsa215 and *E. minutum* Pei191) and other endosymbiotic bacteria. Accession numbers for references genomes used in this analysis are provided in Supplementary Table 4. Note that the genes marked with a “*” or a “!” were not in the final assemblies, but their reads were detected by either Megan (!) or by read mapping onto the same gene from another closely related organisms (*) or both (!*) (see Supplementary Figure 2 as an example).

**Supplementary Figure 5.**
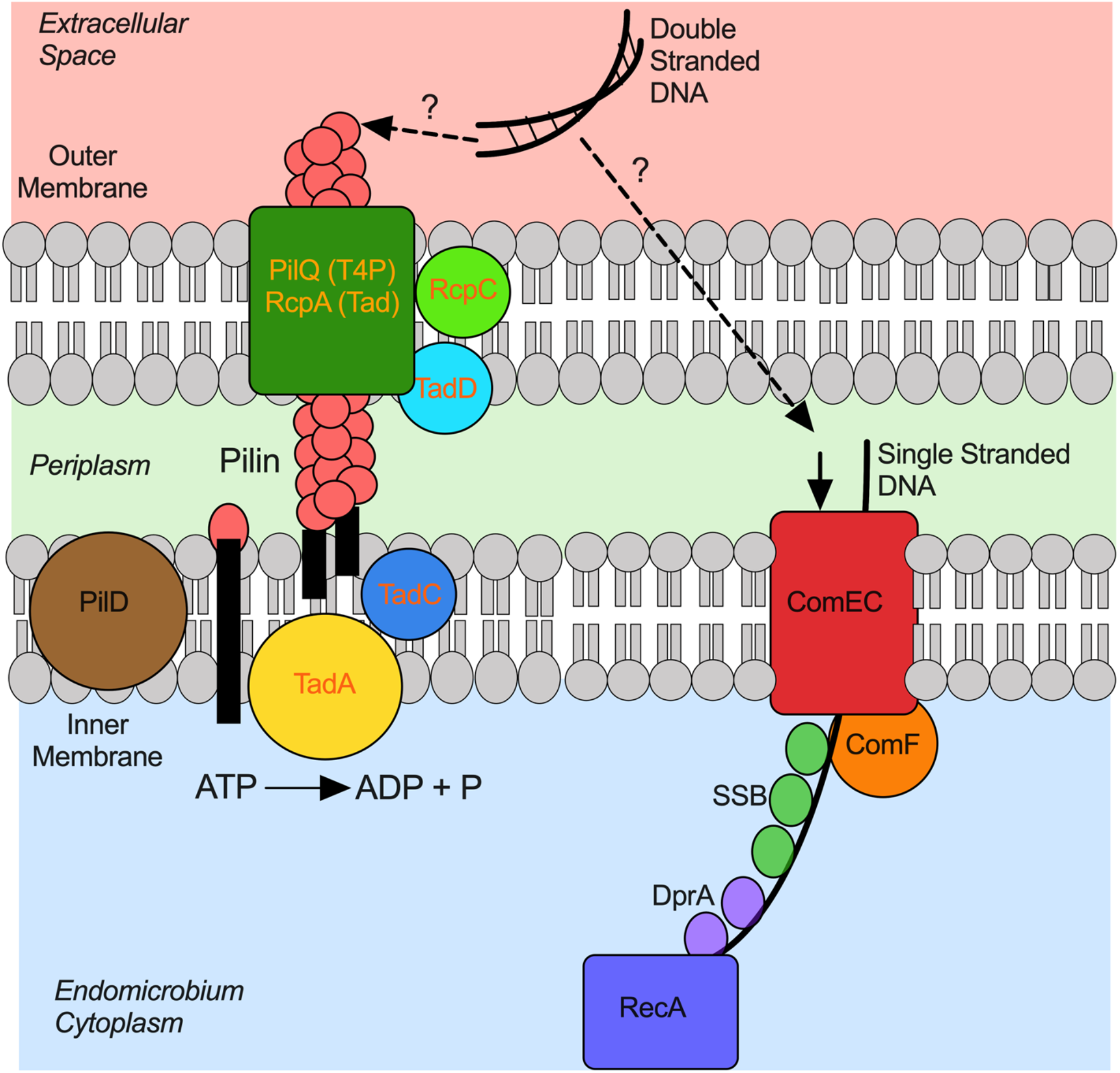
Graphical summary of a putative competence pathway and proteins involved in pilus assembly in *Endomicrobium* species. Proteins shared by all *Endomicrobium* species are in black font while those that are only retained in ‘*Ca*. E. agilae’ are colored in orange. All *Endomicrobium* possessed the pre-pilin peptidase (PilD) and one or more genes that encode pilins. ‘*Ca*. E. agilae’ possessed a near-complete *tad* locus/ T2SS as well the T4P secretin (PilQ).

---

Supplementary File S1 spreadsheet contains the following:

Sup. Table 1 Meta-Data

Sup. Table 2 Filtering ref.

Sup. Table 3 Read Numbers

Sup. Table 4 Genome refs.

Sup. Table 5 RPKM values

Sup. Table 6 RPKM charts

Sup. Table 7 Type IV tad genes

Sup. Table 8 Biosynthetic genes

Sup. Table 9 Repair_Compet genes

